# Diverse RNA interference strategies in early-branching metazoans

**DOI:** 10.1101/284349

**Authors:** Andrew D. Calcino, Selene L. Fernandez-Valverde, Ryan J. Taft, Bernard M. Degnan

## Abstract

**Background:** Micro RNAs (miRNAs) and piwi interacting RNAs (piRNAs), along with the more ancient eukaryotic endogenous small interfering RNAs (endo-siRNAs) constitute the principal components of the RNA interference (RNAi) repertoire of most animals. RNAi in non-bilaterians – sponges, ctenophores, placozoans and cnidarians - appears to be more diverse than that of bilaterians, and includes structurally variable miRNAs in sponges, an enormous number of piRNAs in cnidarians and the absence of miRNAs in ctenophores and placozoans.

**Results:** Here we identify thousands of endo-siRNAs and piRNAs from the sponge *Amphimedon queens-landica*, the ctenophore *Mnemiopsis leidyi* and the cnidarian *Nematostella vectensis* using a computational approach that clusters mapped small RNA sequences and annotates each cluster based on the read length and relative abundance of the constituent reads. This approach was validated on 11 small RNA libraries in *Drosophila melanogaster*, demonstrating the successful annotation of RNAi associated loci with properties consistent with previous reports. In the non-bilaterians we uncover seven new miRNAs from *Amphimedon* and four from *Nematostella* as well as sub-populations of candidate cis-natural antisense transcript (cis-NAT) endo-siRNAs. In the ctenophore, the absence of miRNAs is confirmed and an abundance of endo-siRNAs is revealed. Analysis of putative piRNA structure suggests that conserved localised secondary structures in primary transcripts may be important for the production of mature piRNAs in *Amphimedon* and *Nematostella*, as is also the case for endo-siRNAs.

**Conclusion:** Together, these findings suggest that the last common ancestor of animals did not have the entrained RNAi system that typifies bilaterians. Instead it appears that bilaterians, cnidarians, ctenophores and sponges express unique repertoires and combinations of miRNAs, piRNAs and endo-siRNAs.

## BACKGROUND

RNA interference (RNAi) evolved prior to the divergence of extant eukaryotic lineages, possibly in response to threats from parasitic double-stranded RNA species such as retroviruses and transposons [1]. In contemporary animals, three independent RNAi systems comprise the bulk of the small RNA (sRNA) repertoire: micro RNAs (miRNAs); Piwi interacting RNAs (piRNAs); and endogenous small interfering RNAs (en-do-siRNAs). Amongst non-bilaterian animals - sponges, cnidarians ctenophores and placozoans - miRNAs appear to be lost in latter two lineages, with placozoans and ctenophores also lacking key miRNA biogenic enzymes [2-4]. The absence of miRNAs in the sister lineages to the animal kingdom, choanoflagellates, and fungi [2,5,6], suggests the miRNA system has been lost or evolved independently multiple times [7]. Nonetheless, animal miRNAs play fundamental roles in cell type differentiation and maintenance, and their emergence and proliferation is linked to the evolution of complex multicellularity [8]. The prevalence of miRNAs in plants and algae [9] lends further support to the hypothesis that miRNAs may be important regulators of multicellular development. However, miRNAs do not appear to be essential for animal multicellularity given they are missing from the morphologically complex basal metazoans, the ctenophores [3,4].

There are some marked differences in the miRNA systems of sponges, cnidarians and bilaterians. In contrast to bilaterians, which express a complex repertoire of miRNAs in somatic tissues [10-16], miRNA expression in cnidarians is consistently dwarfed by piRNAs [2,17-19]. The miRNAs of the cnidarian *Nematostella vectensis*, while capable of bilaterian-like silencing [20], also regularly silence their targets through extensive base pairing followed by cleavage, as observed in plants, rather than by transcript destabilisation or translational inhibition [19]. The miRNA repertoire in sponges is substantially lower than in cnidarians and bilaterians with only eight, eleven and nineteen currently reported from the demosponges *Amphimedon queenslandica, Stylissa carteri* and *Xestospongia testudinaria* respectively [2,21]. In *Amphimedon*, these differ from other metazoan miRNAs in having a peculiar plant-like pre-miRNA secondary structure, and have no discernable homology with any non-sponge or eumetazoan miRNA [2,22-25].

While questions about miRNA evolution in animals remain unresolved, the roles of the endo-siRNA and piRNA systems during the emergence of the Metazoa have received far less attention. Having evolved deep in eukaryotic evolution, the endo-siRNA pathway was inherited by the crown Metazoa [26,27]. Incontrast, piRNAs appear to be a metazoan innovation [2]. A functional PIWI-piRNA pathway is present in *Hydra* and *Nematostella* [28,29]. This system appears to differ between basal metazoans, with it being absent in placozoans and expressed at relatively high levels in cnidarians [2]. piRNAs have not been studied in ctenophores and the complex repertoire of either endo-siRNAs and piRNAs in non-bilaterian species has yet to be fully documented.

Given the apparent diversity of RNAi systems amongst representatives of early-branching metazoan phyletic lineages, we developed an in silico approach to detail the sRNA components in representatives of these lineages with small RNA libraries and assembled genomes (i.e., *Amphimedon, Nematostella* and the ctenophore *Mnemiopsis leidyi*). We first confirmed the efficacy and accuracy of this approach on 11 well-annotated developmental small RNA libraries from *Drosophila melanogaster*. When applied to the non-bilaterians, this approach identified novel miRNAs, piRNAs and endo-siRNAs and revealed that *Amphimedon, Mnemiopsis* and *Nematostella* have markedly different RNAi repertoires from each other and from bilaterians.

## RESULTS

### The uniformity index as a tool for discriminating RNAi classes

To investigate the sRNA repertoires of *Amphimedon, Mnemiopsis* and *Nematostella*, we developed a method for the annotation of putative precursor transcripts of endo-siRNAs, piRNAs and miRNAs based on Illumina sequenced small RNA libraries (see Methods). This method leverages on the fact that the biogenesis of miRNAs reliably produces sRNAs of a predictable length and sequence [30].

Variation around the most abundant reads within a cluster of a miRNA loci is limited, leading to large numbers of sRNA reads exhibiting low sequence diversity. In contrast, without the guidance of binding partners involved in miRNA production, Dicer cleaves dsRNA with less discrimination, producing endo-siRNAs of a regular length, typically 21-22 nucleotides (nts), but with far greater sequence variability [31-39]. As a consequence, endo-siRNAs loci typically generate a higher diversity of sRNAs that are lower in relative abundance compared to miRNA loci. Likewise, piRNA biogenesis involves limited specificity over the 5’ and 3’ ends produced by the catalytic components of the pathway, resulting in a highly diverse population of piRNAs generally 26-30 nt in length arising from each loci [40-44].

The uniformity of sRNA reads comprising a given cluster can be measured by what we term the uniformity index - the ratio of the total abundance of sRNA reads comprising a cluster (counts) and the number of distinct sRNA reads from that same cluster. For example, a miRNA-like hairpin comprised of 16 counts but only three distinct reads results in a uniformity index of (16/3) or 5.3 while an endo-siRNA like hairpin comprised of 16 counts comprising 12 distinct reads results in a uniformity index of (16/12) or 1.3 (Additional file 1). Calculating this index for each sRNA cluster enables segregation of high uniformity (HU) clusters (such as miRNAs) from low uniformity (LU) endo-siRNA and piRNA clusters, as we demonstrate in *Drosophila*. Amongst the segregated HU clusters are repetitive sequences as well as miRNAs and other biologically significant sRNA clusters which can be secondarily annotated.

Developmental small RNA libraries from *Nematostella* [19] and *Amphimedon* and two replicate small RNA libraries from *Mnemiopsis* [45], were included in our analysis. In addition to the non-bilaterian datasets, we analysed eleven developmental small RNASeq libraries from *Drosophila* [46]. As the sRNA repertoire of *Drosophila* is well characterised, we first determined if the classification pipeline produced results consistent with prior published analyses [43,46-48].

### Discrimination and annotation of RNAi classes in Drosophila

*Drosophila* is one of the most well-annotated and widely studied model organisms in terms of its small RNA repertoire. Of the three RNAi classes, are the best annotated. In total there are 258 miRNAs currently deposited in miRBase (release 21) and 150 of these have been annotated with high confidence [49]. We were able to identify 139 previously reported miRNAs clusters (54% of total) including 121 high confidence miRNAs (81% of total; Additional files 2, 3). The UI of miRNA clusters averaged 122.5 compared to 1.8 for endo-siRNA clusters. No new miRNA candidates were identified in *Drosophila*.

The piRNA repository piRBase currently details over 28 million individual piRNA sequences in *Drosophila* [50]. From the 11 *Drosophila* datasets examined here, we identified 8,929 putative piRNA clusters. Of these, 8,915 (99.8%) overlap a previously reported piRNA sequence (Additional file 3).

In *Drosophila*, endo-siRNAs are less well annotated than either miRNAs or piRNAs. As no central endo-siRNA database has yet been established, we produced a reference database of endo-siRNA loci from those reported in six previous publications (Additional file 3) [32-35,51,52]. This reference database comprises 1210 clusters spanning ∼5.7 million base pairs (bp) or 3.3% of the *Drosophila* genome. Our analysis identified 3,517 endo-siRNA clusters covering approximately 1.4% of the *Drosophila* genome (Additional file 3). An intersection of our reference dataset based on previous publications with the newly identified endo-siRNA cluster loci identified 13.3% congruence (467 loci) between the two. This represents a significant enrichment compared to what would be expected if the reference dataset and the newly identified endo-siRNA clusters had uncorrelated genomic distributions (*p* <0.00001; see Supplementary Methods, Additional file 4). 86.7% of the endo-siRNA clusters identified by our pipeline were not found in the reference. This may be due to the incompleteness of the limited reference endo-siRNA dataset.

Evidence of a ping-pong biogenesis signature (a bias for a uridine at position one and an adenosine at position 10) [40,41,43] was found in the putative piRNAs from both the *Drosophila* adult female and adult male body libraries as well one of the 2 - 4 day old pupal libraries (Additional file 5). Such a signature was not found in any of the putative endo-siRNAs in which, as expected, only a position one-uridine bias was observed (Additional file 6) [31,37].

To confirm an association between transposons and the putative endo-siRNA and piRNA clusters, the genomic positions of all clusters were intersected with those of annotated coding sequences, including exons, introns, 5’ untranslated regions (5’ UTRs) and 3’ untranslated regions (3’ UTRs), and known and unknown transposons (based on sequence similarity to Repbase entries). Clusters that did not overlap with these genomic elements were deemed to be ‘intergenic’. As anticipated, multi-mapping endo-siRNAs and piRNAs derive primarily from transposons (Fig. 1) [53,54]. In addition, we found that unique endo-siRNA clusters frequently map to exons, 5’ and 3’ UTRs in coding genes (Fig. 1), with unique endo-siRNA clusters underrepresented in introns suggesting that endo-siRNA production occurs after intron splicing.

**Figure 1.**
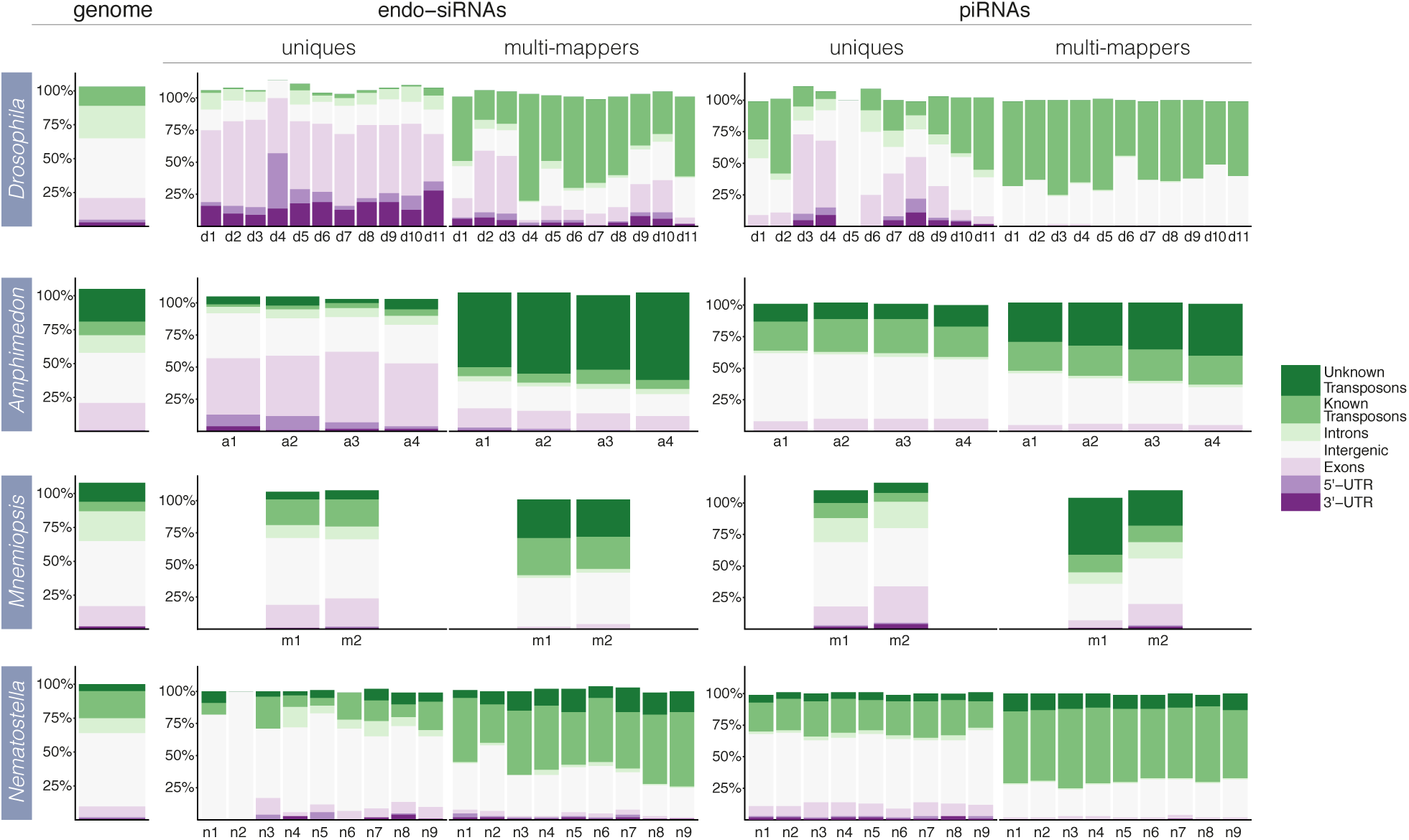
Genomic context of endo-siRNA and piRNA cluster expression for unique and multi-mapping clusters. Each colour-coded segment represents the percentage of endo-siRNA or piRNA clusters mapping to the specified genomic elements. Percentages slightly exceed 100% due to some regions of the genome encoding multiple types of element. The genome column shows the percentage of the genome covered by the specified genomic elements. For *Drosophila*, d1) 12-24 hour embryo, d2) first instar larvae 1, d3) first instar larvae 2, d4) third instar larvae 1, d5) third instar larvae 2, d6) 0-1 day pupae, d7) 2-4 day pupae 1, d8) 2-4 day pupae 2, d9) male adult body, d10) female adult body, d11) female adult head; *Amphimedon* a1) pre-competent larvae, a2) competent larvae, a3) juvenile, a4) adult; *Mnemiopsis*, m1) *Mnemiopsis* 1, m2) *Mnemiopsis* 2; *Nematostella*, n1) unfertilized eggs, n2) blastula, n3) gastrula, n4) early planula larvae, n5) late planula larvae, n6) metamorphosing, n7) primary polyp, n8) male adult, n9) female adult.

The program Randfold [55] was used to test the likelihood that the secondary structures predicted to form from the precursor transcripts of endo-siRNA and pi-RNA clusters could occur by chance. Briefly, Randfold compares the minimum free energy of the predicted secondary structure of a native sequence to the minimum free energies of randomised versions of itself. For each library, Randfold scores were generated for endo-siRNA and piRNA clusters and these were compared to all other clusters (i.e. all clusters other than those under investigation) from the libraries in question (Fig. 2). Both unique and multi-mapping endo-siRNA clusters in *Drosophila* show evidence of secondary structure while putative piRNA transcripts do not (Fig. 2). This is consistent with most models of endo-siRNA and piRNA biogenesis in bilaterians in which some endo-siRNAs are cleaved from secondarily structured primary transcripts while piRNAs are not [53]. Given that the putative piRNA and endo-siRNA clusters identified here have proven to be consistent with previously reported properties, we deemed our method to be satisfactory for naive identification and annotation.

**Figure 2.**
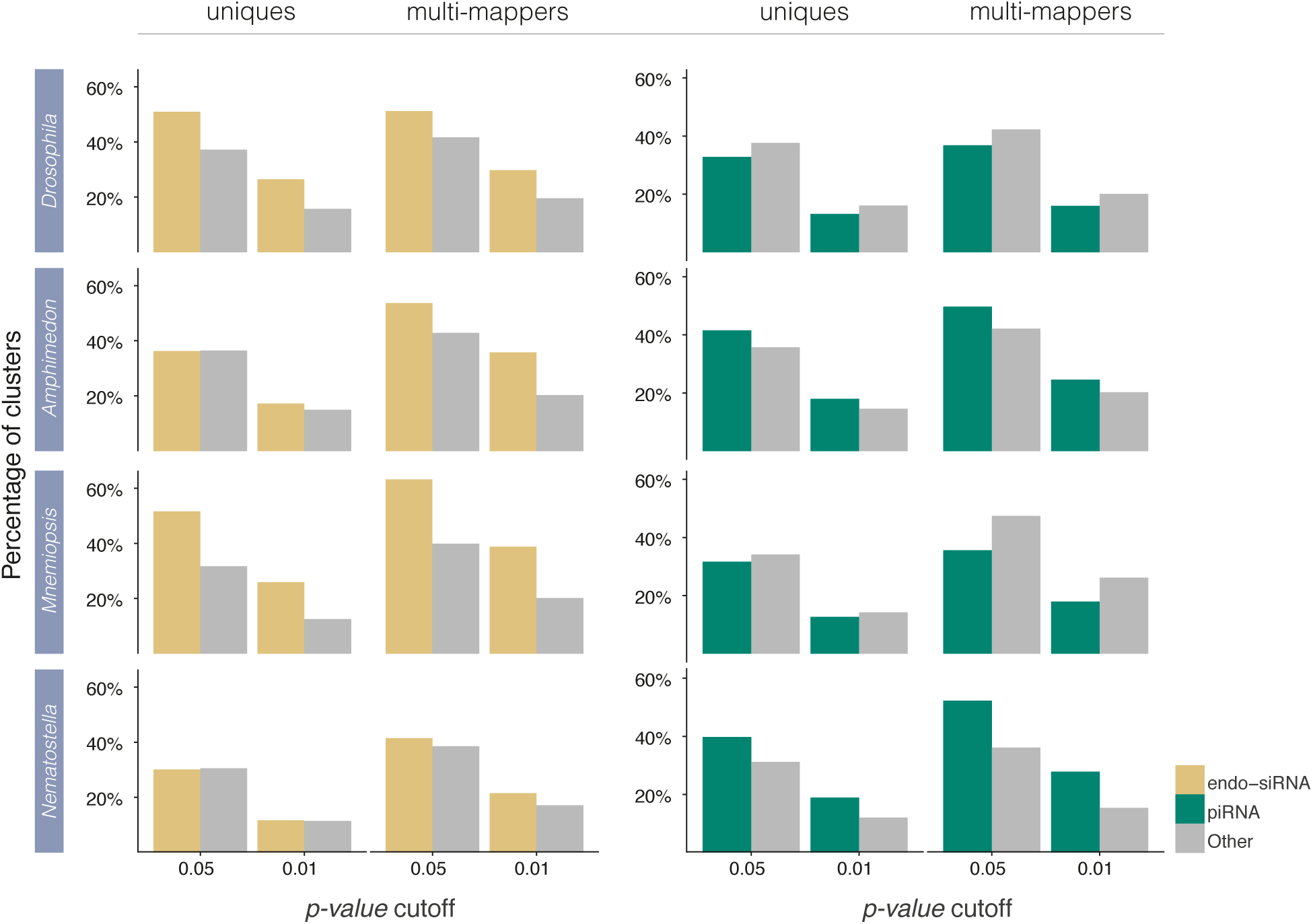
Randfold results for endo-siRNA and piRNA clusters. Each bar represents the percentage of clusters with Randfold p-values equal to or less than the values stated on the X-axis. The more significant the p-value cutoff, the more confidence there is that the secondary structure of the native sequence is more stable than a randomised version of itself. For each graph, the Randfold scores of either endo-siRNAs or piRNAs are compared to the Randfold scores of all clusters not annotated as endo-siRNAs or piRNAs. For each species, all available datasets were pooled.

### Discrimination and annotation of RNAi classes in non-bilaterians

Using the same approach undertaken in *Drosophila*, we surveyed the miRNA, piRNA and endo-siRNA repertoire of *Amphimedon, Nematostella* and *Mnemiopsis*. The numbers of clusters corresponding to each RNAi class in each species are summarised in Table 1.

**Table 1:**
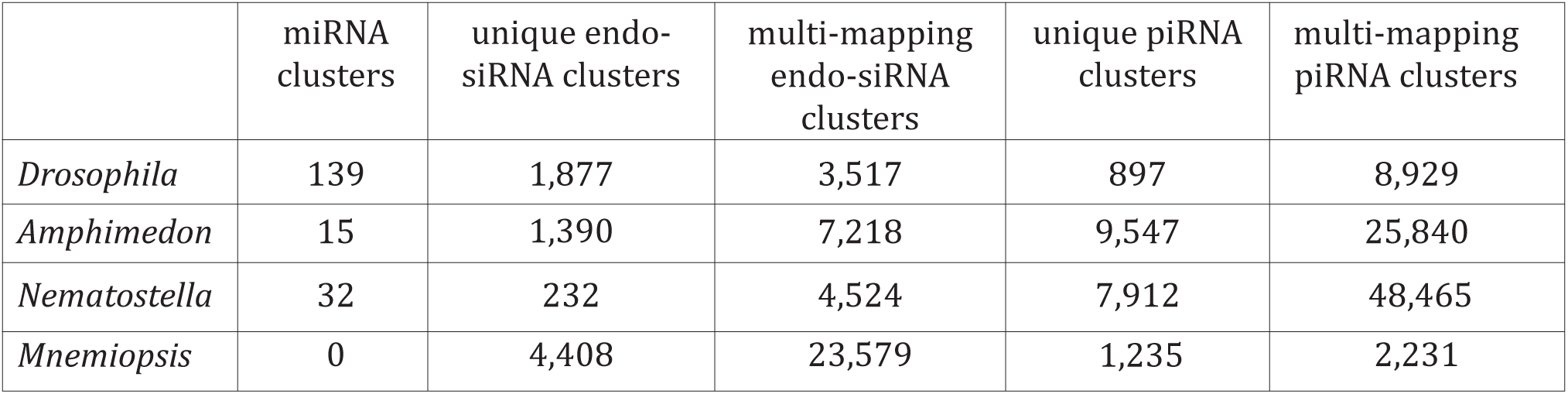
Number of annotated miRNA, piRNA and endo-siRNA clusters in Drosophila, Amphimedon, Nematostella and Mnemiopsis.

Our analysis identified all eight previously reported miRNAs from *Amphimedon*, 62 of the previously reported 141 miRNAs from *Nematostella* and confirmed the absence of miRNAs in *Mnemiopsis*. In addition, we identified seven new miRNA candidates from the sponge including a second copy of aqu-miR-2016 located just over 1 kilobase (kb) from the originally annotated copy, and four new miRNAs in the cnidarian, all of which are copies of previously reported miRNAs (Additional files 7, 8, 9). None of the newly identified sponge miRNAs share sequence similarity with the miRNAs of *Stylissa* and *Xestospongia* [21].

Structurally, three of the new miRNA hairpins(aqu-miR-temp-1,4,6) resemble canonical metazoan pre-miRNAs while the remaining three (aqu-mir-temp-2,3,5) are more similar to the eight previously described long-form miRNAs in *Amphimedon* (Additional file 7) [2]. All of these candidates possess either low numbers of reads mapping to their passenger strands or variable passenger strand 5’ ends [56], however as these characteristics are present in some high confidence miRNAs, such as human hsa-miR-126 [56], we annotate these six loci as candidate novel miRNAs. The remaining HU endo-siRNA-like clusters consist of a mixture of snoRNA, tRNA and rDNA loci, and clusters with highly multi-mapping dominant reads, endogenous hairpin RNAs (hp-RNA; Additional file 10) [57] and secondary structures not consistent with any known sRNA class. Unlike in *Drosophila* where evidence of a ping-pong biogenesis signature was only found in two of the 11 libraries, a bias for a 5’ uridine and an adenosine at position 10 was detected in all *Amphimedon* and *Nematostella* libraries and one of the two *Mnemiopsis* libraries (Additional file 5). As expected, endo-siRNA clusters only exhibit a bias for a 5’ uridine (Additional file 6) [31,37].

As in *Drosophila*, unique *Amphimedon* endo-siRNA clusters frequently map to coding genes (Fig. 1). In contrast, distributions of unique endo-siRNAs do not show a bias towards coding genes in *Mnemiopsis* or *Nematostella*. Unique endo-siRNAs in these species map to coding genes with a frequency more similar to that which would be expected if they were randomly distributed throughout the genome (Fig. 1). In all species, multi-mapping endo-siRNA and piRNA clusters tend to map to transposons.

Randfold analysis of unique endo-siRNA clusters in *Amphimedon* and *Nematostella* show that they are no more likely to form secondary structures than the putative transcripts of all other unique clusters (i.e. all identified sRNA clusters not including endo-siRNAs). In contrast, unique endo-siRNA clusters in *Mnemiopsis*, as in *Drosophila*, show evidence of secondary structuring, as do multi-mapping endo-siRNA clusters in all four species (Fig. 2).

Unexpectedly, putative piRNA transcripts of *Amphimedon* and *Nematostella* show evidence of secondary structure for both unique and multi-mapping clusters while those in *Mnemiopsis* are more similar to the unstructured piRNAs known from bilaterians (Fig. 2).

### Variation in overall RNAi complements in basal metazoans

The relative contributions of sRNAs differ amongst the representative sponge, ctenophore, cnidarian and bilaterian species (Fig. 3). In *Amphimedon* and in all but one *Drosophila* developmental stage, miRNAs comprise the bulk of mapped sRNAs while endo-siRNAs and pi-RNAs are dominant in *Mnemiopsis* and *Nematostella* respectively. Except for the *Nematostella* libraries, a substantial proportion of each library remains unassigned to one of the three RNAi classes (Fig. 3). This is likely due to the stringent requirements set here for annotating sRNA clusters (see Supplementary Methods, Additional file 4) and the presence of non-RNAi related sRNAs produced by each animal.

**Figure 3.**
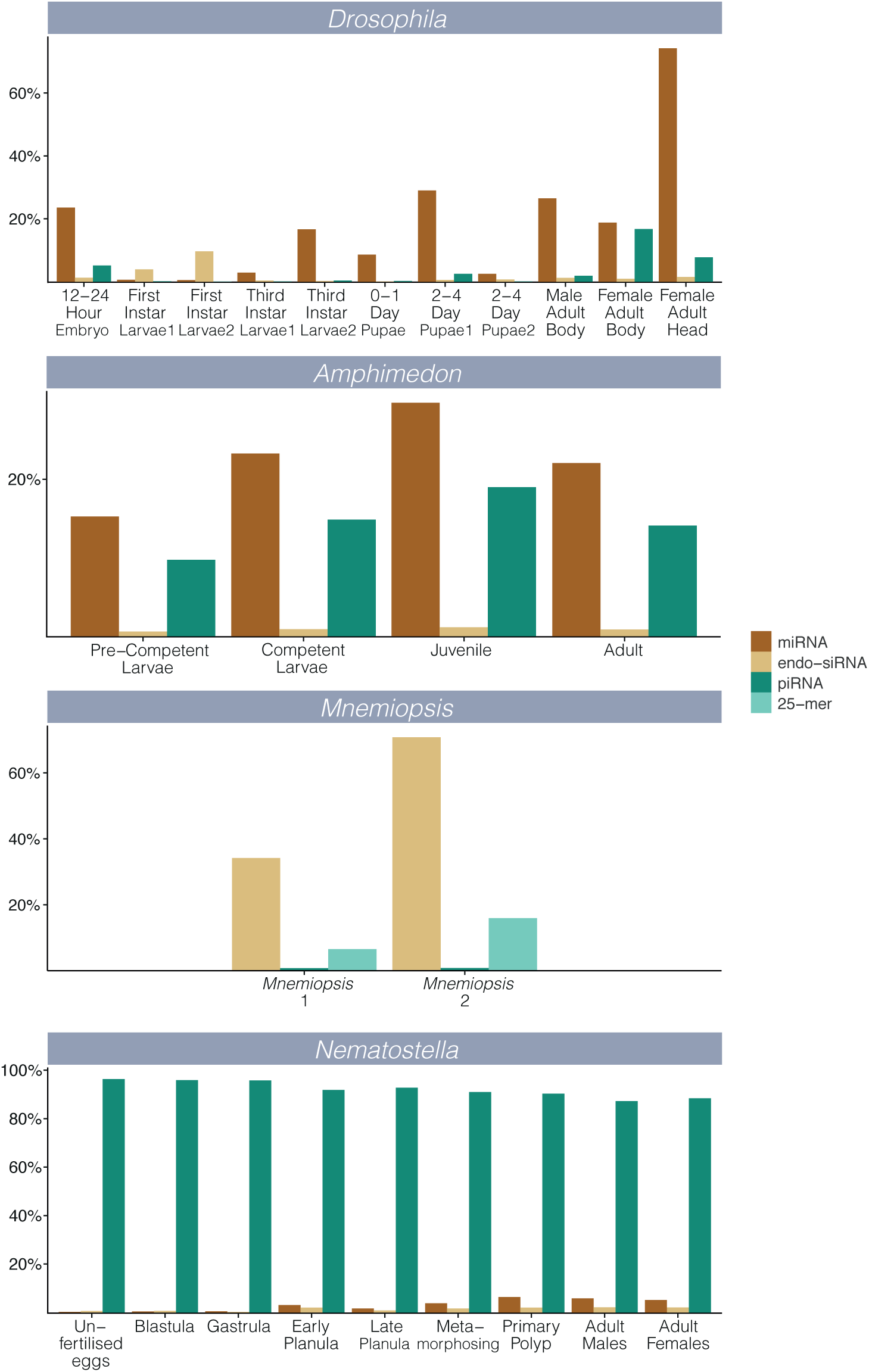
Library contributions from each RNAi component as a percentage of total library depth. Total contributions of miRNAs, endo-siRNAs, piRNAs and *Mnemiopsis* 25-mer clusters to total library depth. For each, only a single copy of each multi-mapping read was considered.

### Developmental dynamics of endo-siRNA and pi-RNA expression

Co-expression of endo-siRNA and piRNA clusters across developmental time was investigated for *Drosophila, Amphimedon* and *Nematostella* (Fig. 4; Additional file 11); *Mnemiopsis* was excluded due the absence of developmental data. This analysis highlights differences in the expression dynamics of endo-siRNAs and piR-NAs; while many endo-siRNAs are co-expressed in the *Nematostella* male adult and female adult libraries, the populations of piRNAs in these two samples appear to be more distinct. Likewise for *Amphimedon*, endo-siR-NA co-expression is highest for the two larval libraries whereas piRNAs appear to be more consistently expressed in all four developmental stages.

**Figure 4.**
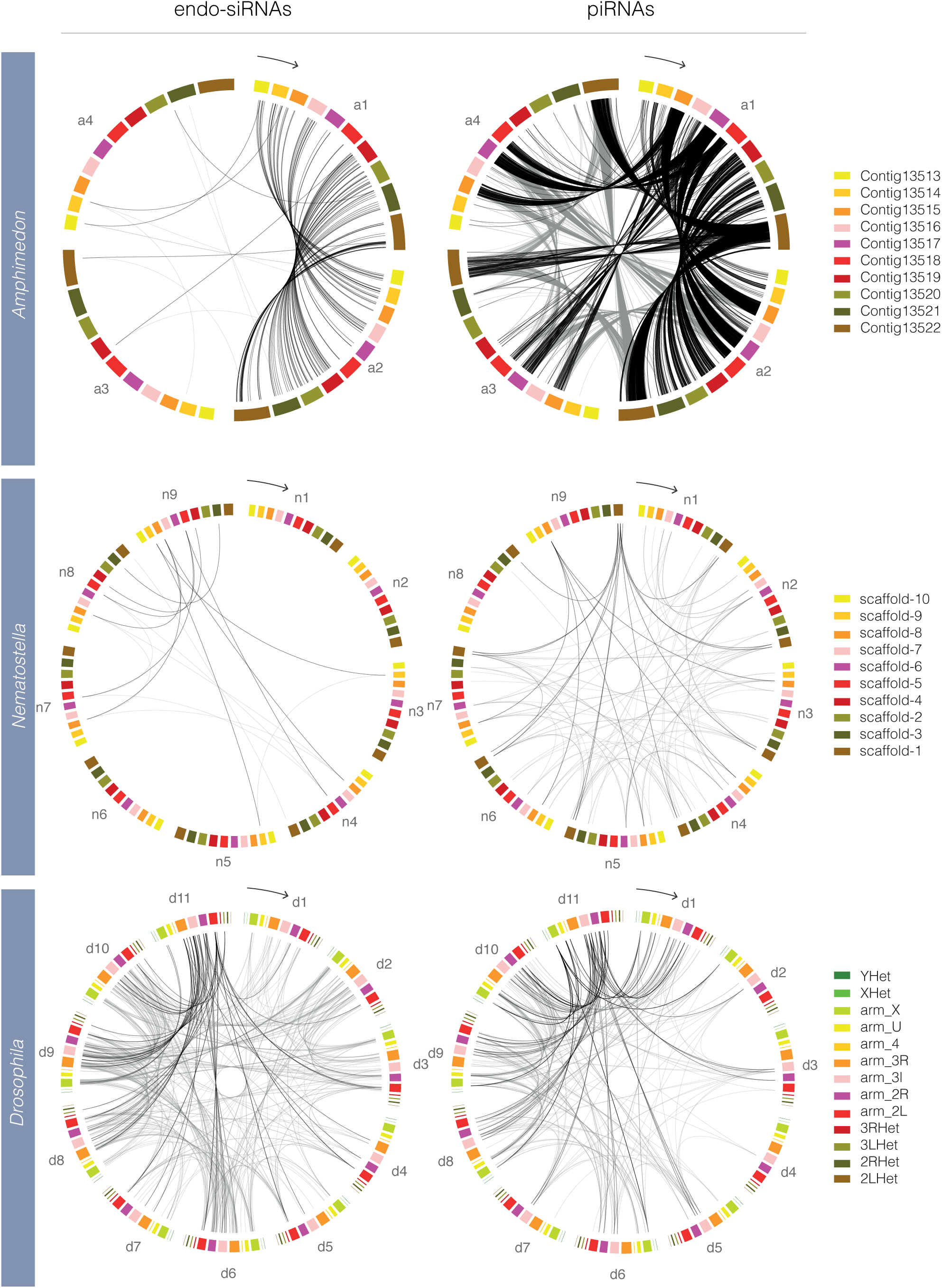
Co-expression of uniquely-mapping endo-siRNA and piRNA clusters. Each plot is divided in to groups of coloured scaffolds/chromosomes, each of which represents a developmental stage; four stages in *Amphimedon*, nine stages in *Nematostella* and 11 stages in *Drosophila*. For each plot, the earliest developmental stage is marked with an arrow indicating the chronological order of developmental stages. Links between scaffolds/chromosomes indicate co-expression from a particular endo-siRNA or piRNA cluster in the two linked developmental stages. For *Drosophila*, all chromosomes are represented while for *Amphimedon* and *Nematostella*, the ten largest genomic scaffolds were used. Beginning with the developmental stage indicated by the arrow, the stages for *Amphimedon, Nematostella* and *Drosophila* are as per Fig. 1. For each species, the links shared with a single developmental stage are coloured black for emphasis while the rest are coloured grey. For *Amphimedon* the emphasised stage is the pre-competent larvae, for *Nematostella* the female adult and for *Drosophila*, the female adult head.

### Mnemiopsis 25-mer cluster annotation

In addition to putative endo-siRNA and piRNA clusters, a substantial proportion of *Mnemiopsis* reads were found to be approximately 25 nt in length (Additional file 12). As it was unknown whether these constituted a separate class of a previously reported sRNA type or a novel sRNA class; *Mnemiopsis* clusters producing ∼25 nt reads (hereafter referred to as 25-mer clusters) were investigated separately. Small RNAs from *Mnemiopsis* 25-mer clusters have a bias for a uridine at their 5’ end, as is common for en-do-siRNAs but no ping-pong biogenesis signature was identified (Additional file 13). No mapping bias to any genic feature including transposons was observed (Additional file 14) and there was no evidence of secondary structure of the putative 25-mer precursor transcripts (Additional file 15). As no evidence of structure or function was identified for 25-mer clusters, further work is required to determine whether they are biologically significant.

## DISCUSSION

RNA interference systems are important post-transcriptional regulators in metazoans. A thorough overview of the repertoire and developmental dynamics of these systems is lacking for most taxa, however, resulting in an incomplete picture of their evolutionary trajectory. We sought to address the early evolution of the metazoan miRNA, piRNA and endo-siRNA systems through the annotation of their small RNA components in the demosponge *Amphimedon*, the ctenophore *Mnemiopsis* and the cnidarian *Nematostella*. We developed a method for the clustering and annotation of mapped sRNA libraries. Application of this method to the bilaterian *Drosophila* recapitulated the results of previous studies [32-35,40,41,51,52], suggesting this approach could be applied to other species.

Specifically, we demonstrate that *Drosophila* miRNAs account for the highest number of mapped reads and piRNAs and endo-siRNAs are dynamically expressed and frequently map to transposons. Endo-siRNAs display a bias for a 5’ uridine and the ping-pong biogenesis signature can be detected in annotated piRNAs. We also showed that 99.8% of annotated *Drosophila* piRNA clusters identified using this method map to previously reported piRNAs and that at least 13% of endo-siRNAs also correspond to previously reported endo-siRNA generating loci. In agreement with the established models of endo-siRNA and piRNA biogenesis, secondary structure appears to be important for *Drosophila* endo-siRNA clusters but not for piRNA clusters.

Using this strategy, we detected all previously reported miRNAs in the *Amphimedon* datasets and 44% of known miRNAs from *Nematostella*. We also confirmed the absence of miRNAs in the ctenophore and showed that endo-siRNAs and piRNAs are the most abundant RNAi classes in *Mnemiopsis* and *Nematostella* respectively. In *Amphimedon*, as in *Drosophila*, unique endo-siRNAs derive primarily from the exons and UTRs of coding genes, consistent with these being derived from mature spliced mRNAs in both species.

Primary transcript secondary structure does not appear to be a requirement for piRNA biogenesis [58], although one study identified a role for the RNA helicase MOV10L1/Armitage in unwinding localised secondary structures of piRNA precursors in mice and *Drosophila* [59]. Orthologues of this helicase can be found in *Amphimedon* (NCBI: XP_019853676.1), *Nematostella* (NCBI: XP_001626596.1, XP_001637169.1) and *Mnemiopsis* (NHGRI: ML005359a). Our analysis did not find any evidence of conserved piRNA cluster secondary structure in *Drosophila* or *Mnemiopsis*, however *Amphimedon* and *Nematostella* piRNA clusters do appear to be structured. This suggests that sites of conserved localised secondary structure within primary piRNA transcripts may be a main source of piRNA production in *Amphimedon* and *Nematostella*.

Unique and multi-mapping endo-siRNA clusters in *Drosophila* and *Mnemiopsis* have a propensity to form secondary structures while only multi-mapping endo-siRNA clusters appear to in *Amphimedon* and *Nematostella*. As endo-siRNA directed RNA interference is most efficient for targets with full-length complementarity [60], most uniquely mapping endo-siRNAs are expected to silence transcripts arising from the antisense strand from which their host gene was transcribed [61]. Consistent with this, Randfold analysis of the predicted secondary structures formed by *Amphimedon* unique endo-siRNA clusters showed that they are more likely to occur by chance than are the secondary structures formed by multi-mapping endo-siRNA clusters.

Given that (i) *Amphimedon* does not encode an RNA dependent RNA polymerase (RdRP), (ii) secondary structure is probably less important for the biogenesis of most unique endo-siRNAs and (iii) the most efficient targets of unique endo-siRNAs are likely found antisense to themselves, it follows that most unique endo-siRNAs are likely to be the products of cis-Natural Antisense Transcripts (cis-NATs) [35,62] rather than hairpin RNAs. Of the 40,122 coding gene models for *Amphimedon* [63], 8,133 are predicted to be cis-NATs. While this only represents 20.3% of the total coding genes, nearly 50% of all coding gene-mapping unique endo-siRNA clusters align to these genes.

Unique endo-siRNA clusters in *Drosophila* also align to coding genes, although both unique and multi-mapping endo-siRNA clusters show evidence of forming secondary structures. Despite this, the 16% of genes that form cis-NAT pairs in this species account for 22% of all mature coding gene-mapping unique endo-siRNA clusters demonstrating that cis-NATs are the likely source of some uniquely mapping endo-siRNAs in *Drosophila*. Differences in the rate of cis-NAT endo-siRNA production observed between *Drosophila* cell types [52] may account for the lower overall rate detected in comparison to *Amphimedon*. The more compact *Amphimedon* genome may also be responsible for a higher rate of overlapping antisense transcripts [63,64].

## CONCLUSIONS

The RNAi repertoires of non-bilaterian metazoans - sponges, ctenophores and cnidarians – differ both from each other and from the canonical RNAi repertoire of bilaterians. Although largely comprised of the same three major systems that constitute the bilaterian RNAi repertoire, the degree to which miRNAs, piRNAs and endo-siRNAs are expressed varies substantially between these basal lineages. The unexpected differences in the RNAi repertoire of bilaterians, cnidarians, ctenophores and sponges uncovered here, suggests that while the last common ancestor of extant animals employed miRNA, piRNA and endo-siRNA systems, these were not integrated into the shared ancestral gene regulatory system. This is in contrast to bilaterians, which in general appear to use a common RNAi system [10-16]. Following the emergence of these major metazoan RNAi pathways, lineage-specific evolutionary trajectories appear to have resulted in divergent RNAi strategies evolving in each basal metazoan lineage.

## METHODS

### Biological sampling and sequencing

Detailed methods can be found in Supplementary Methods. Briefly, *Amphimedon* material was collected from Heron Island, Australia and RNA extracted using Tri Reagent (Sigma Aldrich). Small RNA libraries were constructed either with the Illumina TruSeq small RNA kit (adult sponge) or the Epicentre ScriptMiner Small RNA-Seq Library Preparation Kit (pre-competent larvae, competent larvae and juvenile). Sequencing was performed on an Illumina HiSeq 2000 at the Institute for Molecular Bioscience, Brisbane, Australia. *Mnemiopsis* (SRS355925, SRS355926) [3], *Nematostella* (SRR039731, SRR039754, SRR039764, SRR039762, SRR039760, SRR039758, SRR039756, SRR039726, SRR039727) [19] and *Drosophila* (SRR013604, SRR018039, SRR016854, SRR013601, SRR013603, GSM360260, SRR013600, SRR013602, GSM360256, GSM360257, SRR014367) [46] datasets were acquired either from NCBI’s Sequenced Read Archive (SRA, http://www.ncbi.nlm.nih.gov/sra/) or from NCBI’s Gene Expression Omnibus (GEO, http://www.ncbi.nlm.nih.gov/geo).

### Small RNA cluster generation and annotation

Small RNA reads were mapped to their respective genomes and clustered using bedCluster.pl [66]. A 150 bp window was defined for cluster generation, reflecting the approximate length of the long pre-miRNAs typical of *Amphimedon* [2] and in recognition that miRNA, piRNA and endo-siRNA biogenesis results in products located in overlapping or close genome proximity to one another, all of which derive from an original primary transcript (or two in the case of natural antisense endo-siRNAs) [38]. Only clusters composed of at least three distinct reads (non-perfectly overlapping) and at least 51 bp in length were considered. Clusters corresponding to previously reported miRNAs were annotated as such. tRNAs were predicted with tRNA-scan-SE [67] and snoRNAs with snoSeeker [68], and clusters mapping to these locations were annotated. The minimum free energy of each cluster was defined using RNALfold [69] and Randfold analysis [55] with 100 randomisations.

### endo-siRNA, piRNA and 25-mer cluster annotation

Both endo-siRNAs and piRNAs clusters were annotated based on the read length composition of their constituent sRNAs. For endo-siRNAs, clusters with peaks of expression at 20, 21 or 22 nt were first selected. If the sum of the reads constituting the peak read length plus or minus one nucleotide was greater than the total number of reads of all other size classes, these were annotated as endo-siRNA clusters. For piRNA annotation an sRNA peak of 26, 27 or 28 nt was required for the non-bilaterians while for *Drosophila*, 24, 25 or 26 nt were selected, reflecting the shorter length of piRNAs in this species [40]. For *Mnemiopsis* 25-mer clusters 24, 25 or 26 nt peak clusters were also selected.

### Circos plots

Circos plots [70] were constructed that describe the co-expression of clusters in different developmental contexts. Links were formed between corresponding genomic loci from two developmental stages if those loci co-expressed either an endo-siRNA or piRNA cluster in both temporal contexts.

## ADDITIONAL FILE INFORMATION

### Additional file 1

#### Figure S1. Demonstration of High Uniformity and Low Uniformity sRNA clusters

Description of data: Two hypothetical hairpin RNAs demonstrating the difference between a high uniformity and a low uniformity clustering. In (a), a total of 16 reads composed of just three distinct reads map to a hairpin RNA giving a uniformity index of 5.3. In (b), 16 reads also map to a hairpin RNA but these are composed of 12 distinct reads resulting in a uniformity index of just 1.3. The high uniformity cluster (a) is composed of an equal number of reads to the low uniformity cluster (b) however these reads are less evenly distributed along the length of the source hairpin RNA.

### Additional file 2

#### Uniformity of Drosophila endo-siRNA and miRNA clusters

Description of data: Endo-siRNA clusters (yellow) display a consistently lower uniformity of small RNA expression (ratio of total read counts:distinct reads) in comparison to miRNA clusters (red) for both unique clusters (above) and multi-mapping clusters (below).

### Additional file 3

#### Locations of annotated RNAi loci from Drosophila, Amphimedon, Nematostella and Mnemiopsis

Description of data: Genomic loci of annotated miRNA, piRNA, endo-siRNA and 25-mer clusters in all four species.

### Additional file 4

#### Supplementary Methods

Description of data: Detailed methods

### Additional file 5

#### Figure S3. Nucleotide biases of piRNA clusters

Description of data: Nucleotide biases along the length of all sRNAs mapping to predicted piRNA clusters. sRNAs were anchored at their 5’ nucleotide and biases are displayed as a percentage the contribution of each nucleotide at each position. Of note is the tendency for a uracil at position 1 and an adenosine at position 10 in most libraries which together comprise the ping-pong piRNA biogenesis signature.

### Additional file 6

#### Figure S4. Nucleotide biases of endo-siRNA clusters

Description of data: Nucleotide biases along the length of all sRNAs mapping to predicted endo-siRNA clusters. sRNAs were anchored at their 5’ nucleotide and biases are displayed as a percentage of the contribution of each nucleotide at each position. Of note is the tendency for a uracil at position 1 which is present in all libraries except the *Drosophila* 1st instar larval libraries.

### Additional file 7

#### Figure S5. New Amphimedon miRNA candidates

Description of data: Wiggle plots and predicted secondary structures of mapped reads across the length of previously described miRNA miR-2016a, the newly identified miR-2016b and six novel miRNA candidates (aqu-mir-temp-1-6). For each cluster, the library with the most mapped reads to each loci was used to construct the graph.

### Additional file 8

#### Figure S6. New Nematostella miRNA candidates

Description of data: Wiggle plots and predicted secondary structures of four newly identified miRNAs in the sea anemone. All four miRNAs are new copies of previously identified miRNAs.

### Additional file 9

#### Table S1. New miRNA data

Description of data: Sequence and genomic location data for the newly identified *Amphimedon* and *Nematostella* miRNAs.

### Additional file 10

#### Figure S7. Amphimedon endogenous hairpin RNAs

Description of data: Wiggle plots and predicted secondary structure of three long highly complementary endo-siRNAs from *Amphimedon* with unevenly distributed mapped sRNA populations.

### Additional file 11

#### Figure S8. Co-expression of multi-mapping endo-siRNA and piRNA clusters across development

Description of data: Each plot is divided into groups of coloured scaffolds/chromosomes, each of which represents a developmental stage. For each plot, the earliest developmental stage is marked with an arrow indicating the chronological order of the following developmental stages. Links between scaffolds/chromosomes indicate co-expression from a particular endo-siRNA or piRNA cluster in the two linked developmental stages. For *Drosophila*, all chromosomes are represented while for *Amphimedon* and *Nematostella*, the ten largest genomic scaffolds were used. Beginning with the developmental stage indicated by the arrow, the stages for *Amphimedon, Nematostella* and *Drosophila* are as per Fig. 1. For each species, the links shared with a single developmental stage are coloured black for emphasis while the rest are coloured grey. For *Amphimedon* the emphasised stage is the pre-competent larvae, for *Nematostella* the female adult and for *Drosophila*, the female adult head.

### Additional file 12

#### Figure S9. Read length distribution of all mapped sRNAs from Mnemiopsis

Description of data: Numbers of distinct reads (red) and total read counts (blue) of all mapped sRNA size classes from the Woods Hole, MA, USA library (A) and the Miami, FL, USA library (B). Of particular note are the peaks of mapped sRNAs at 21 and 25 nt in both libraries.

### Additional file 13

#### Figure S10. Nucleotide biases of Mnemiopsis 25-mer clusters

Description of data: Nucleotide biases along the length of all sRNAs mapping to 25-mer clusters. sRNAs were anchored at their 5’ nucleotide and biases are displayed as a percentage of the contribution of each nucleotide at each position. Of note is the tendency for a uracil at position 1.

### Additional file 14

#### Figure S11. Genomic context of 25-mer cluster expression from Mnemiopsis

Description of data: Each colour-coded segment represents the percentage of 25-mer clusters mapping to the specified genomic elements. Percentages slightly exceed 100% due to some regions of the genome encoding multiple types of element. The genome column demonstrates the percentage of the genome covered by the specified genomic elements. Of note is the lack of enrichment of 25-mer clusters from coding genes or transposons.

### Additional file 15

#### Figure S12. Randfold results for Mnemiopsis 25-mer clusters

Description of data: Each bar represents the percentage of clusters with Randfold p-values equal to or less than the values stated on the X-axis. The more significant the p-value cutoff, the more confidence there is that the secondary structure of the native sequence is more stable than a randomised version of itself. For each graph, the Randfold scores of either endo-siRNAs or piRNAs are compared to the Randfold scores of all other clusters. Both datasets were pooled for this analysis.

## Ethics approval and consent to participate

No specific ethics approval was required for this project.

## Consent for publication

Not applicable.

## Authors’ contributions

ADC contributed to the design of the project, analysis and interpretation of the data, collected the required material from the field, conducted the laboratory procedures and drafted the manuscript. SLFV, RJT and BMD contributed to the design of the project, analysis and interpretation of the data and drafted the manuscript. All authors have read and approved the final manuscript.

## Acknowledgements

We thank Kelin Ru and and Anupma Choudhary for their assistance and advice with the production of the *Amphimedon* RNASeq libraries and to Paulina Tapia for her assistance with document design. We also thank Carmel McDougall for feedback on the manuscript.

