## Supplementary figures and images for "Diverse RNA interference strategies in early-branching metazoans"

### Supplementary Materials

*Figure S1*

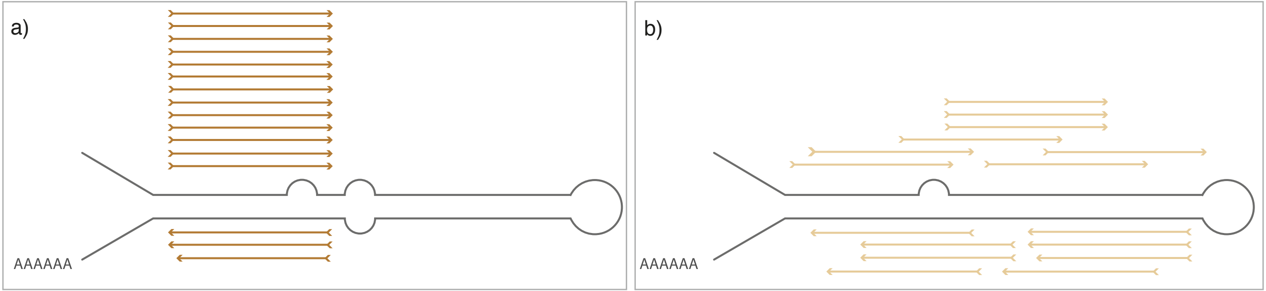

### Supplementary Materials

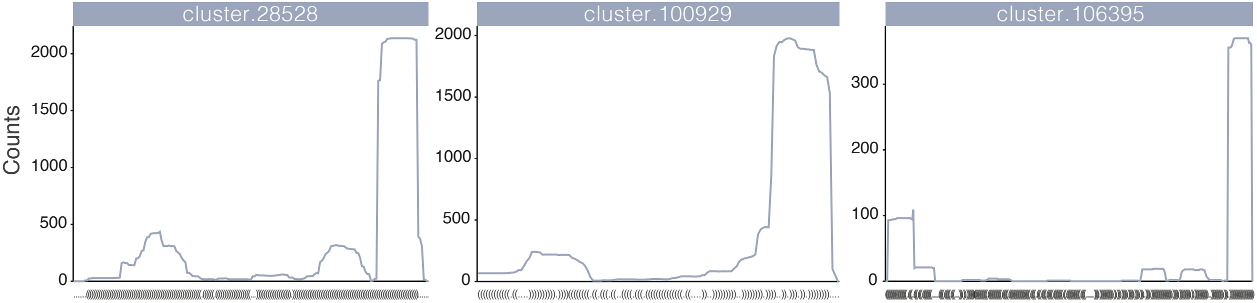

### Supplementary Materials

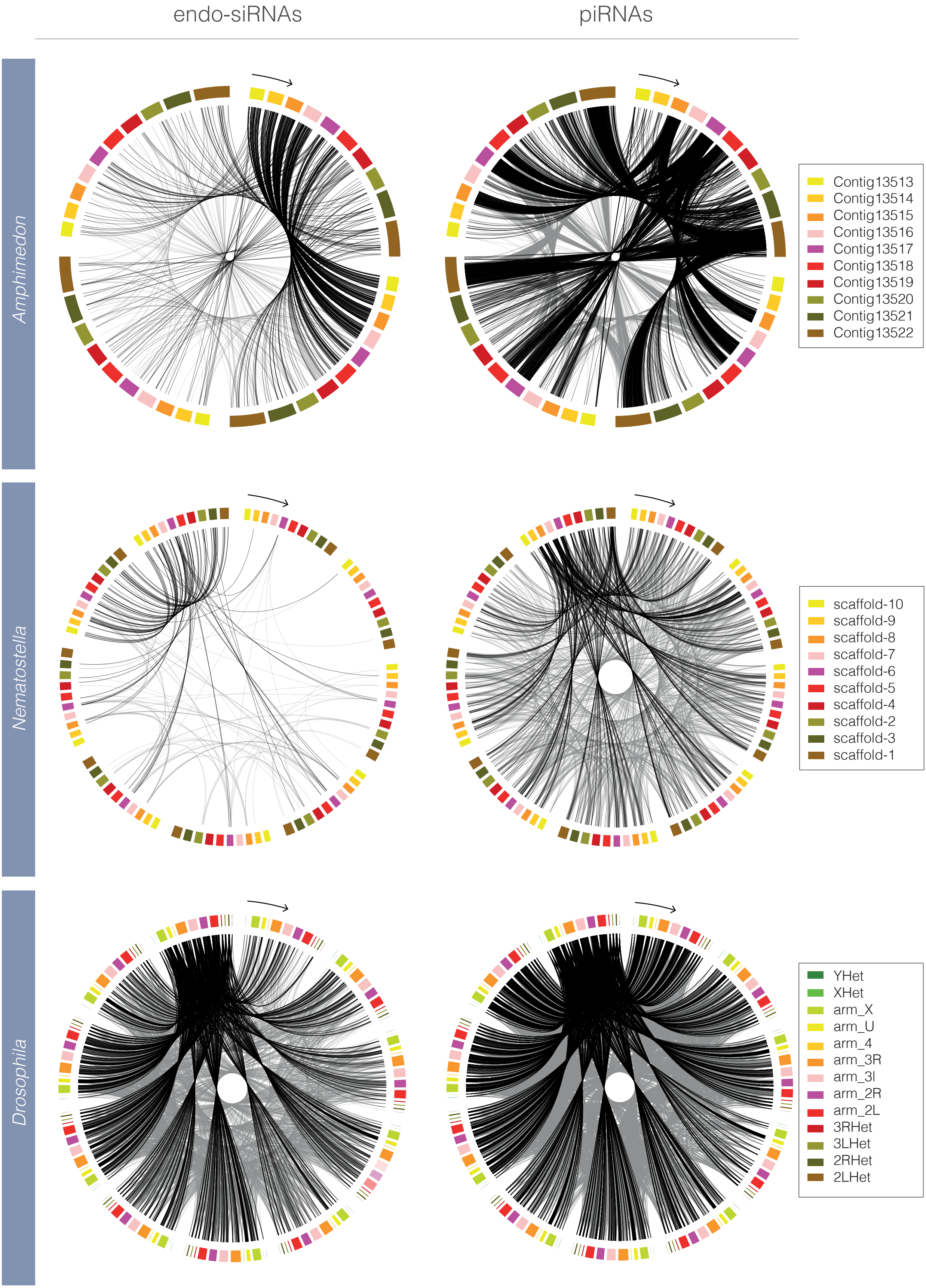

### Supplementary Materials

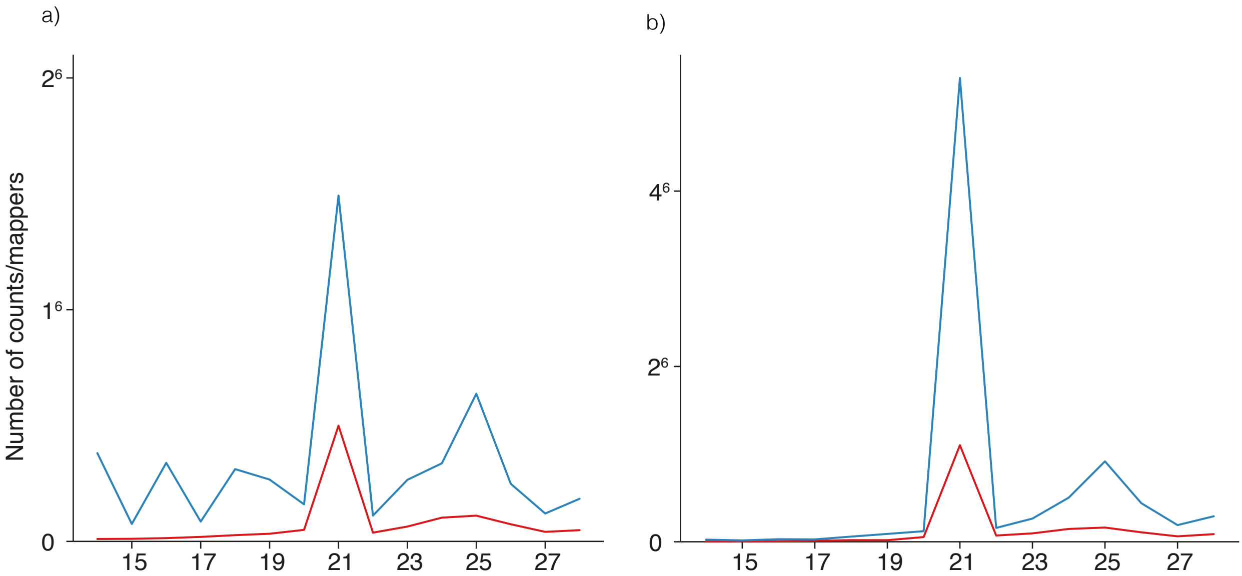

### Supplementary Materials

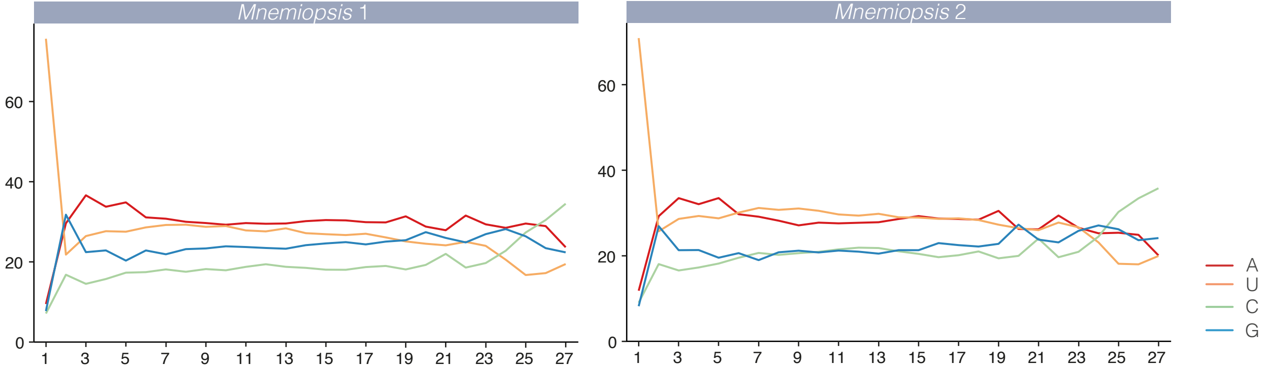

### Supplementary Materials

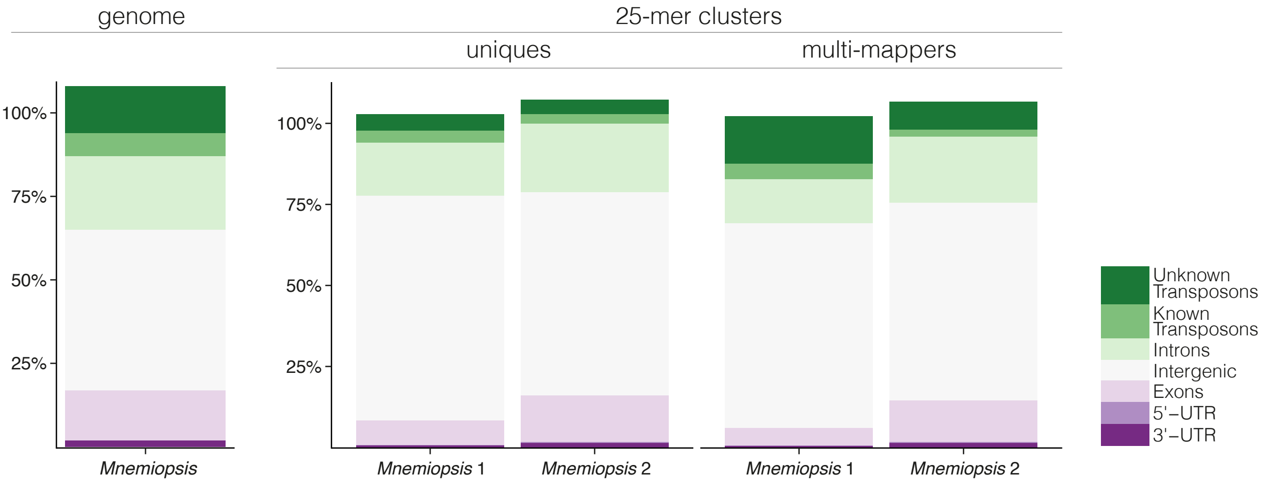

### Supplementary Materials

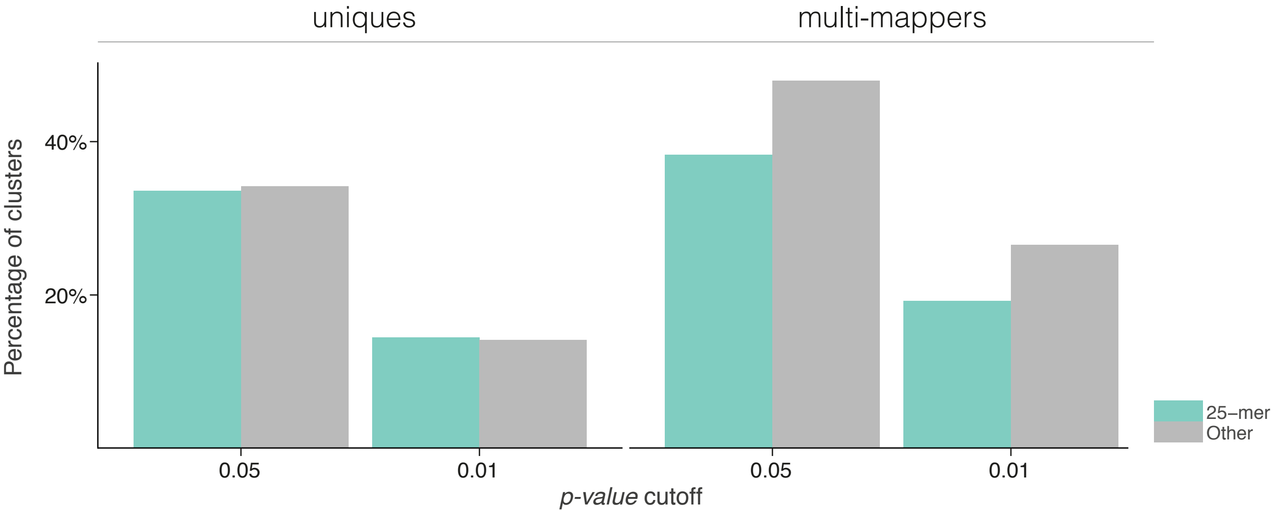

### Supplementary Materials

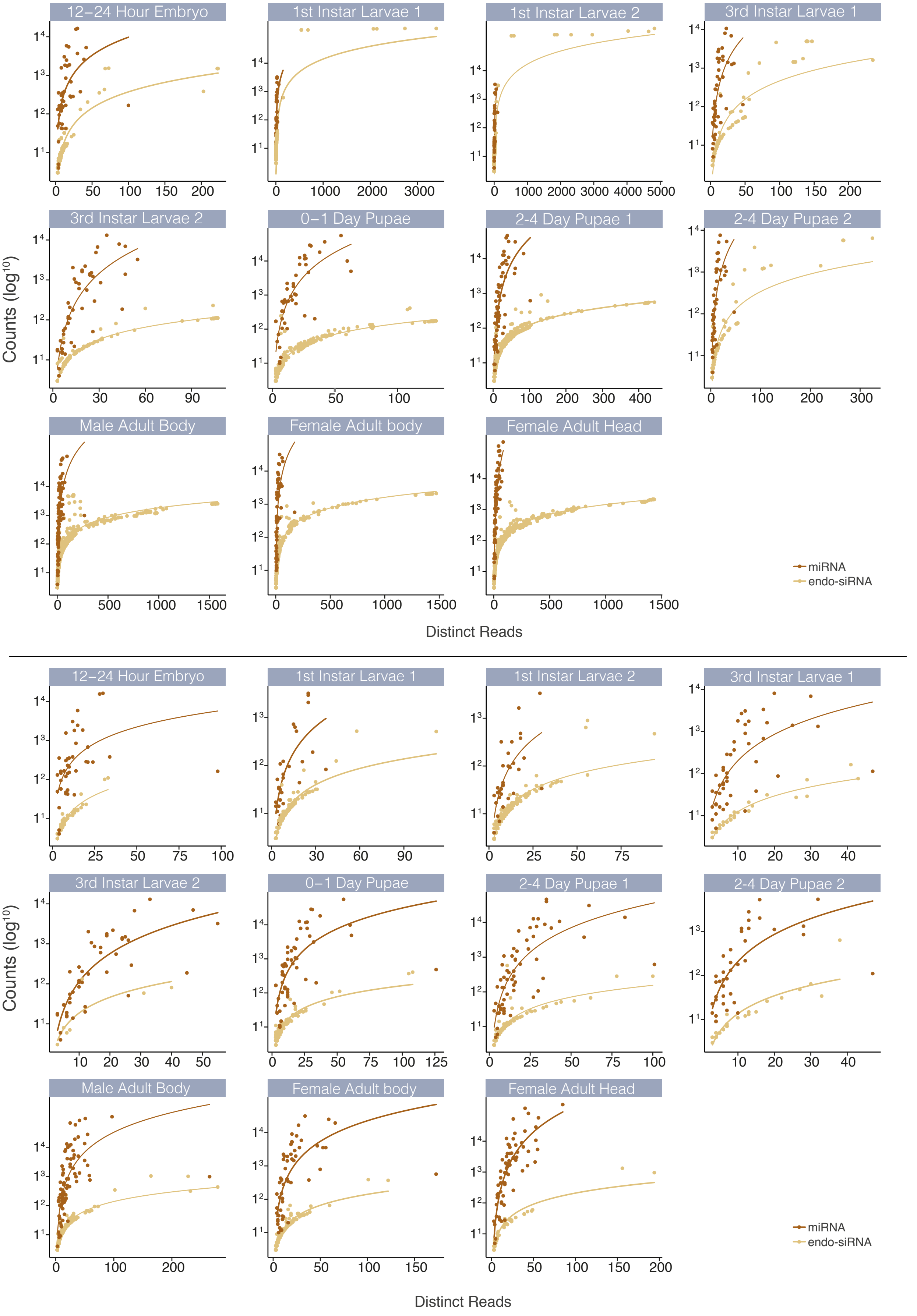

### Supplementary Materials

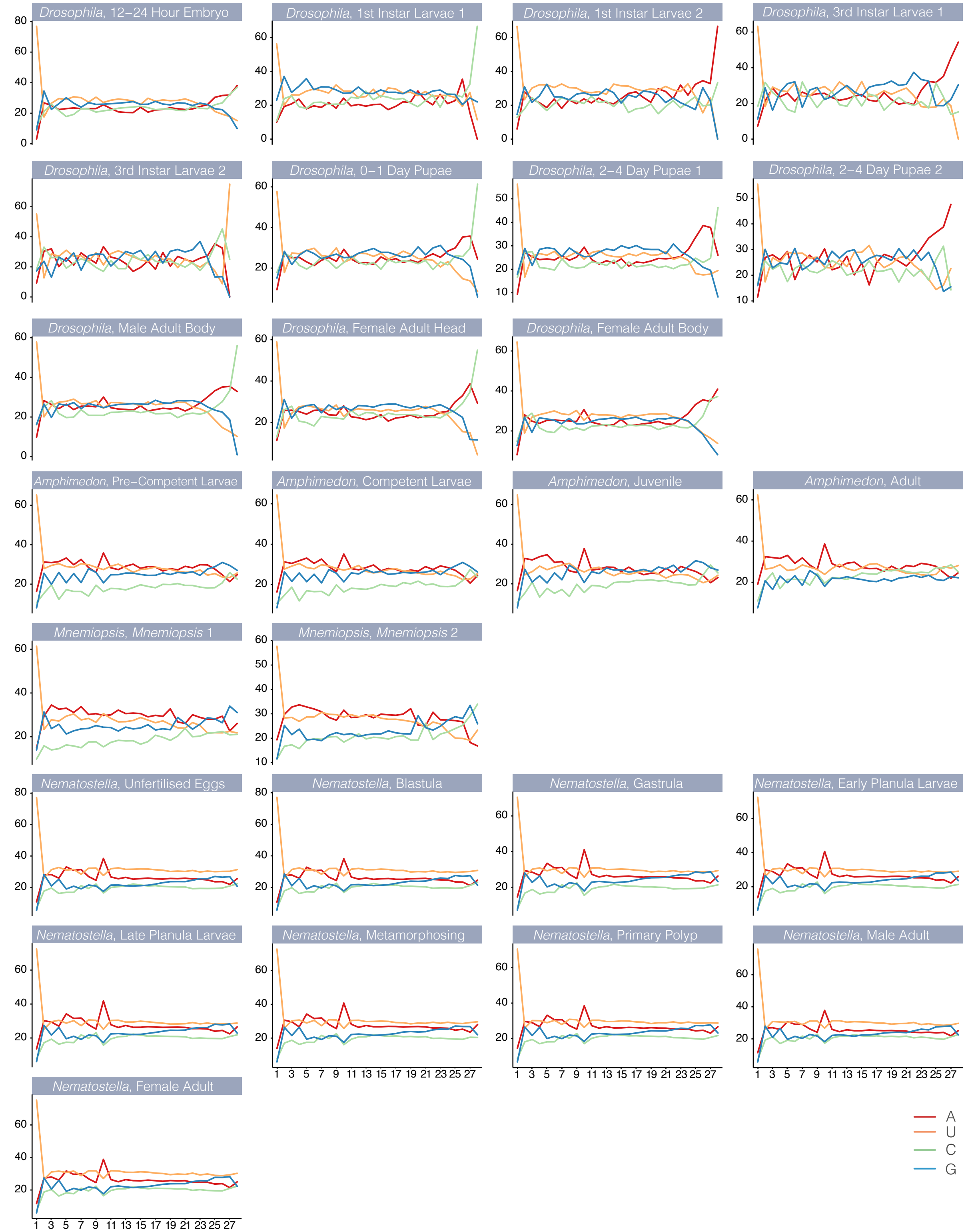

### Supplementary Materials

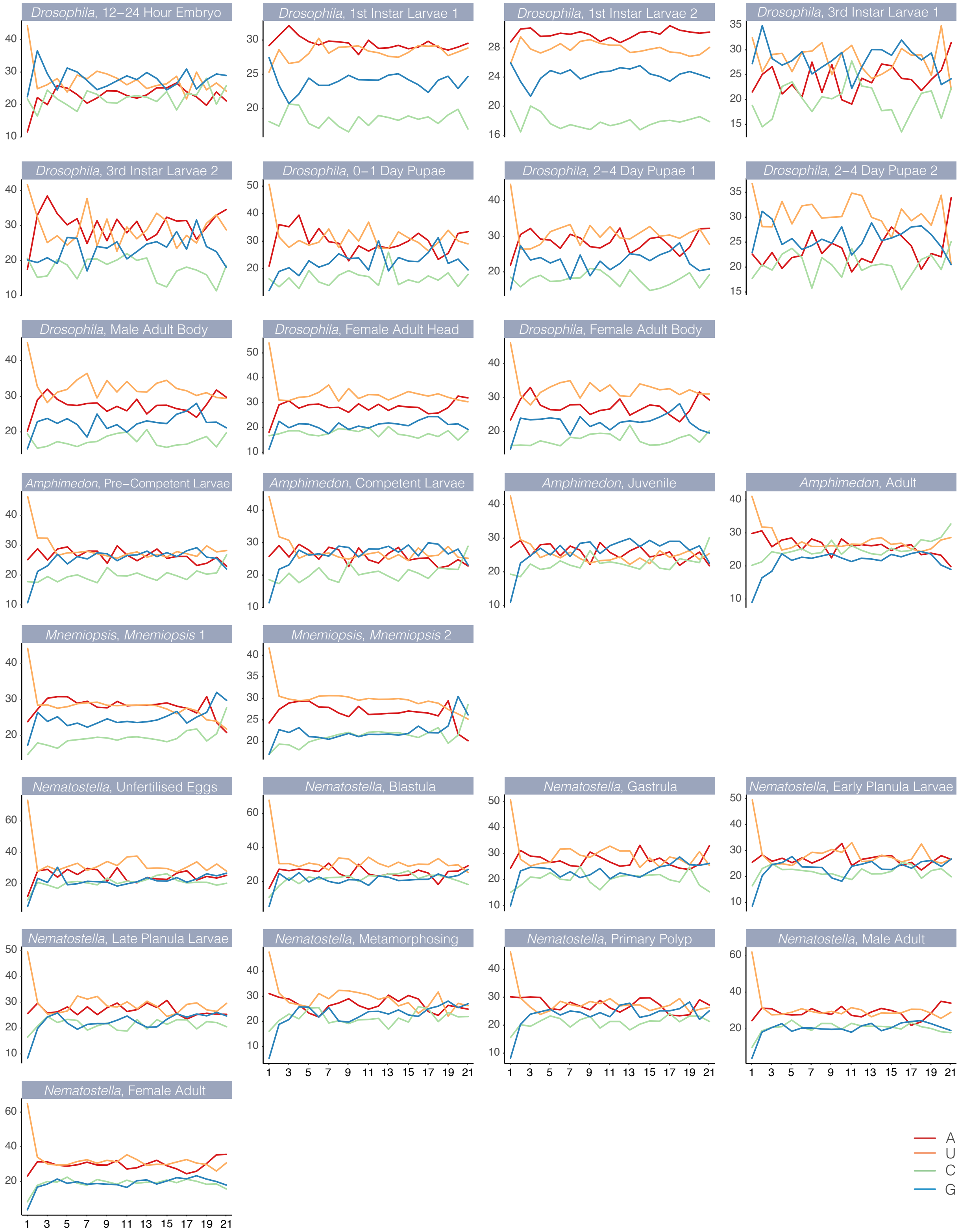

### Supplementary Materials

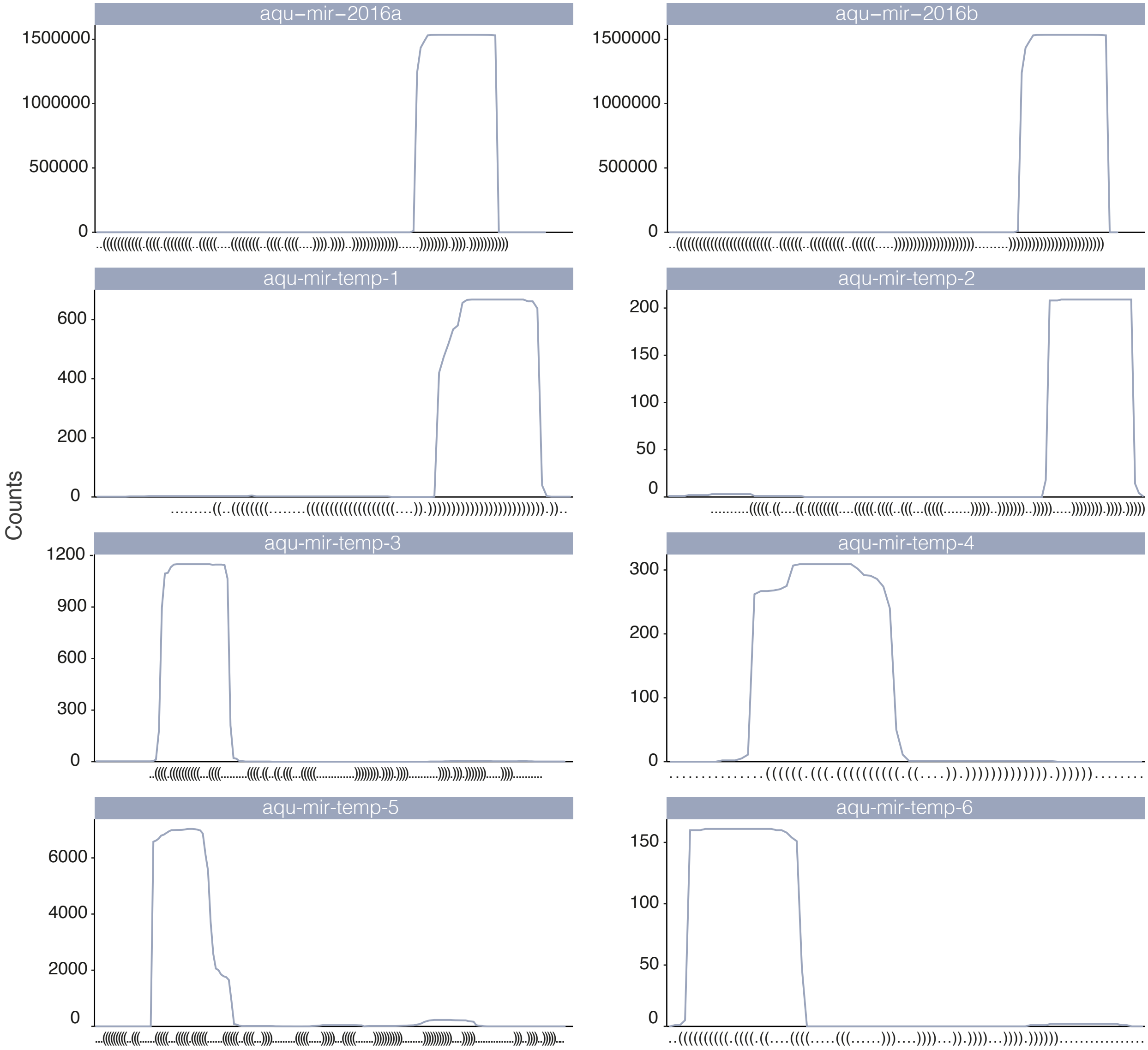

### Supplementary Materials

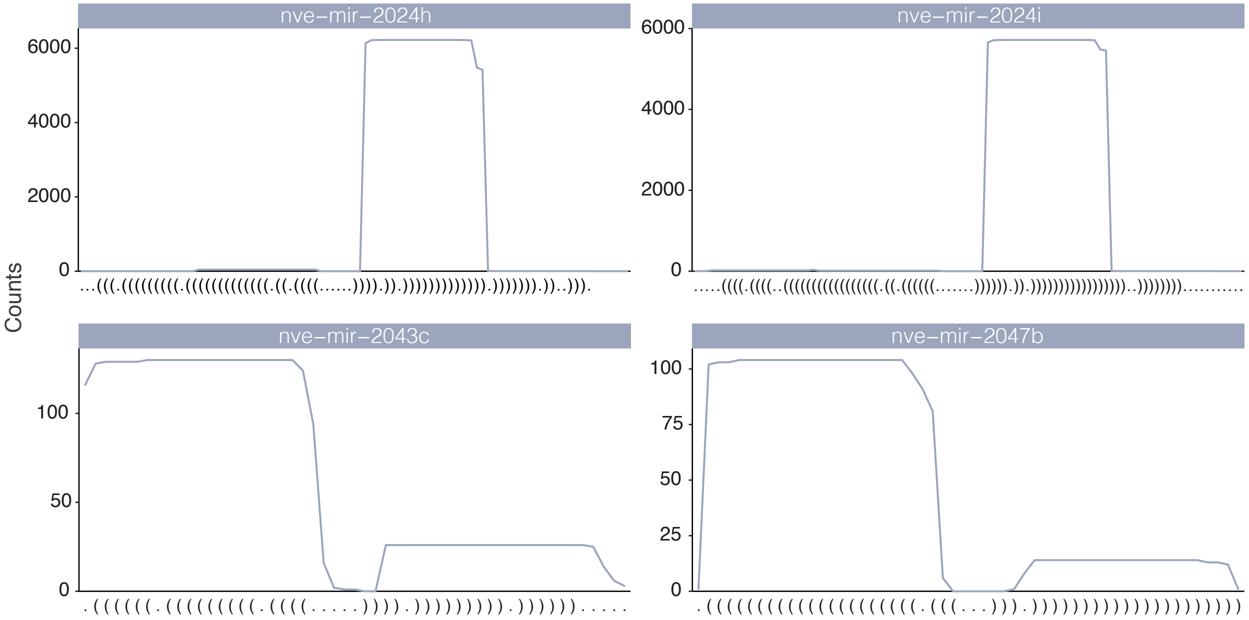
