## Supplementary Materials for "Diverse RNA interference strategies in early-branching metazoans"

### SUPPLEMENTARY METHODS

#### Biological sampling of *Amphimedon queenslandica*

Biological sampling was conducted at the Heron Island Research Station on Heron Island Reef on the Great Barrier Reef, Australia. An adult *Amphimedon* tissue biopsy was taken while snorkelling from the reef flat at Shark Bay. This sample was brought back to the laboratory and immediately transferred to RNALater for cold storage at -20° C until RNA extraction could be conducted. To obtain developmental material, another adult sponge was collected from the reef and brought back to the research station where it was maintained in a tank in flow through seawater continuously pumped form the reef [1]. This sponge was allowed to spawn naturally and at midday, all larvae that were released earlier were cleared from the tank to ensure the age of sampled larvae. After two hours, 0 - 2 hours post-emergence (0-2 hpe) pre-competent larvae [2,3] were sampled and transferred immediately to RNALater for cold storage. The remaining larvae were induced to metamorphose with the introduction of a coralline algae settlement cue [3]. Approximately 10 hours after the introduction of the settlement cue, those larvae that had settled were gently detached and transferred to glass coverslips that were submerged under filtered seawater (FSW) in 24-well plates [4]. Detached individuals rapidly reattach to the new surface whereupon they continue to metamorphose. The 24-well plates were kept in the dark and FSW was changed daily. Three days after re-settlement, the establishment of a contiguous pinacoderm and a single exhalent osculum marked the completion of metamorphosis [1]. These juveniles were sampled and transferred to RNALater for cold storage. In total, between 30 and 50 pre-competent larvae, competent larvae and juveniles were sampled. RNA extraction was conducted with Tri Reagent (Sigma Aldrich) as per the manufacturer's instructions with the following modifications. RNALater was removed from each sample and replaced with at least 500µl of Tri Reagent. Cell disruption was conducted by five to six rounds of heating samples to 55°C for three minutes followed by vortexing for 5-10 seconds. 1-bromo-3-chloropropane was used for phase separation. RNA precipitates were resuspended in 20µl of ultrapure RNAse/DNAse free water.

**Small RNA library preparation and sequencing**

The adult small RNA library was prepared with the Illumina TruSeq Small RNA Sequencing Kit as per the manufacturer's instructions. Pre-competent larval, competent larval and juvenile small RNA libraries were prepared with the Epicentre ScriptMiner Small RNA-Seq Library Preparation Kit as per the manufacturers instructions. Libraries were indexed and the four libraries were pooled with eight others unrelated samples giving a total of 12 samples. These were then split and sequenced over four lanes on an Illumina HiSeq 2000 at the Institute for Molecular Bioscience, Brisbane, Australia.

**Quality control and library mapping**

Output fastQ files from Illumina sequencing were checked for quality with FastQC. 3' adaptor sequences were removed with fastx_clipper from the FASTX-Toolkit (v0.0.13). Collapsed reads were mapped to their respective genomes with bowtie (v0.12) [5] allowing for up to 51 mappings per read but no mismatches between the read and the genome. Those reads that were mapped to the genome 51 times were then removed from the library, leaving only reads that mapped between 1 and 50 times. A second file was produced from those reads that only mapped to a single genomic location.

**Cluster generation**

All sRNAs that map to annotated rRNAs were first removed from all libraries. The remaining reads were clustered using bedCluster.pl with a 150 bp window specified [6]. These clusters were defined as being composed of reads from a single strand or from both strands. Only clusters composed of at least three distinct reads and that were at least 51 bp long were considered further.

Cluster Minimum Free Energy Each cluster was subjected to secondary structure analysis with RNALfold from the Vienna RNA package (v2.05) [7]. For those clusters comprised of reads from both strands, both strands were submitted to RNALfold with the strand that produced the lowest minimum free energy (MFE) retained. If both strands produced equal MFEs, a strand was selected arbitrarily. 30 To assess the likelihood that the structures predicted by RNALfold could have arisen by chance, each was submitted to Randfold [8] and randomised 100 times. Randfold measures the MFE of these randomisations and compares the results to the MFE of the native sequence. The result is a *p-value* assigned to each cluster that describes the likelihood that the native sequence of that cluster will fold to form a secondary structure that is more stable than a randomised version of itself. This can be interpreted as the likelihood that the secondary structure predicted for a cluster has not occurred by chance and thus is likely functionally important.

**Annotation of endo-siRNA, piRNA and 25-mer clusters**

Annotation of endo-siRNA, piRNA and 25-mer clusters was based on properties of the read length distribution of the constituent sRNAs. For endo-siRNAs, clusters with peaks of sRNA expression at 20, 21 or 22 nt were first selected, reflecting the typical length of Dicer cleavage products. Secondly, clusters in which the sum of the reads constituting the peak read length plus or minus one nucleotide was greater than the total number of reads of all other size classes were selected. For piRNA annotation, a similar method was employed but with a sRNA peak requirement of 26, 27 or 28 nt except in the case of *Drosophila* where 24, 25 or 26 nt peaking clusters were selected due to the shorter length of piRNAs in this species [9]. For *Mnemiopsis* 25-mer clusters, like *Drosophila* piRNAs, 24, 25 and 26 nt peaking clusters were selected.

**Analysis of endo-siRNA cluster overlap with reference database**

To test whether the identified endo-siRNA loci overlapped previously reported endo-siRNA loci more often than would be expected by chance, the locations of the endo-siRNA clusters were randomised 100,000 times (randomBed, BEDTools v2.25.0) [10] and then intersected with the reference dataset using overlapSelect [11]. This showed a distribution of overlapping clusters with a mean of 119.5, a standard deviation of 11.6 and maximum and minimum values of 173 and 75 (Fig. 1).

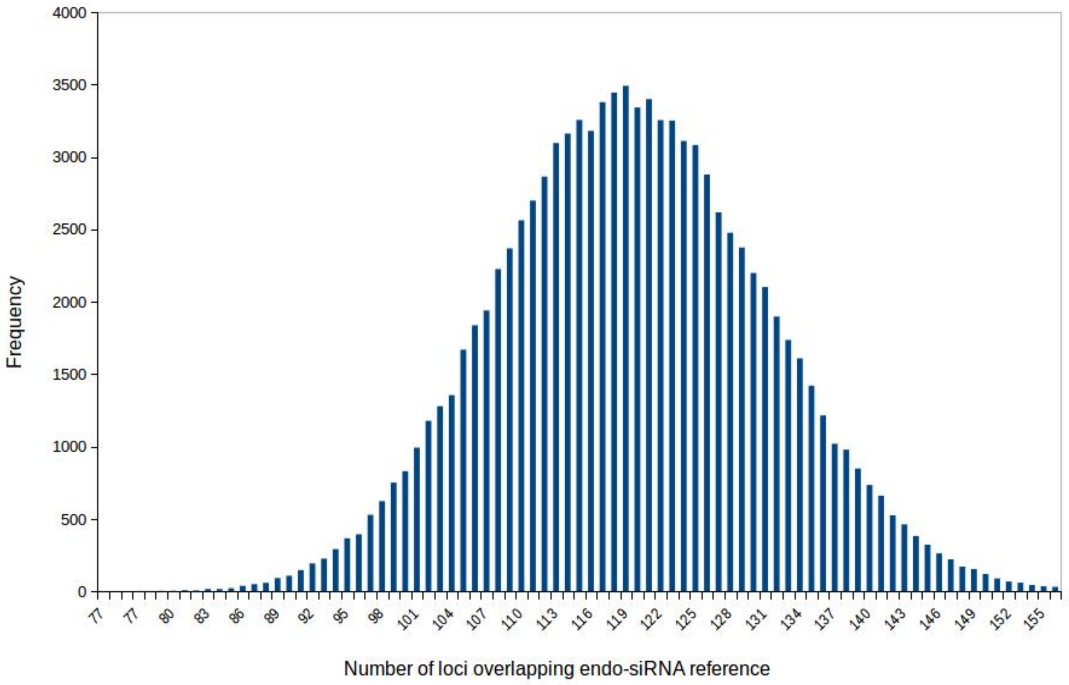


**Fig. 1 Frequency distribution of intersections of endo-siRNA reference database with randomised endo-siRNA loci.**

**Genomic context**

The four genomes were annotated according to their coverage by transposons or coding genes before being intersected with sRNA clusters. RepeatModeler and RepeatMasker [12] were used to identify transposons in all four genomes both with (known) and without (unknown) homology to those in RepBase [13]. Exons, introns, 5’ UTRs and 3’ UTRs were obtained from publicly available sources. Exons, introns, 5’ UTRs and 3’ UTRs that overlapped with predicted transposons were removed. All elements were mapped to the genome with GenomeCoverageBed from the BEDTools package (v2.5.0) [10].

The genomic context of endo-siRNA and piRNA clusters were assessed using overlapSelect from UCSC [11] to determine which elements clusters aligned to. At least 51% of the length of a cluster was required to overlap with a particular feature, otherwise it was deemed to be intergenic. To determine if any HU clusters derive from tRNAs or snoRNAs, HU cluster were intersected with tRNAs as predicted with tRNAScan-SE [14] and snoRNAs as predicted by snoSeeker [15].

**cis-NAT prediction of gene models**

Gene models for the four species were overlapped with themselves using UCSCs overlapSelect [11]. Gene models from opposing strands were aligned to one another and those overlapping another by at least one nucleotide were considered to be cis-NAT genes.

8. Bonnet E, Wuyts J, Rouzé P, Van de Peer Y. Evidence that microRNA precursors, unlike other non-coding RNAs, have lower folding free energies than random

sequences. Bioinformatics. 2004;20:2911–7.
